# Engineered heat-stable variants of *Trypanosoma cruzi* flagellar protein Tc24 enable serological detection of Chagas disease

**DOI:** 10.1101/2025.08.23.671862

**Authors:** Sagar Batra, Christopher Waugh, Gemma Harris, Frederick W. Muskett, Tiberiu-Marius Gianga, Rohanah Hussain, Giuliano Siligardi, Laurence Turpin, Mallory Draye, Pascal Mertens, Barrie Rooney, Michael Miles, Tapan Bhattacharyya, Ivan Campeotto

## Abstract

Chagas disease, caused by the protozoan *Trypanosoma cruzi*, is one of the major neglected diseases globally, killing 12,000 people per year and silently infecting an estimated 7 million people worldwide. Current diagnostic methods are limited by cost, complexity and cold-chain requirements. The *T. cruzi* flagellar protein Tc24 is a promising antigen for serological tests but suffers of poor solubility and heat stability. Here, we computationally engineered and produced three variants of Tc24, which exhibit remarkable heat stability, up to 69°C, and express with higher solubility in *E. coli* compared to the wild-type protein, reducing production costs, eliminating the need for a cold chain and therefore facilitating cost-effective production and storage without refrigeration and even in lyophilised form. These variants remained remarkably stable for 70 days in solution at 25°C and successfully detected antibodies in human sera samples from Chagas disease patients from Northern and Southern regions of Latin America, demonstrating the preservation of their antigenicity. The best-performing engineered variant was incorporated into a prototype of lateral flow test, demonstrating potential for rapid, affordable and accessible Chagas disease diagnostics in resource-limited settings.

## INTRODUCTION

Chagas disease is a chronic life-threatening disease, caused by the parasite *Trypanosoma cruzi* endemic in Latin American countries and represents a significant global health burden worldwide. Chagas disease causes about 12,000 deaths annually, with 7 million people estimated to be infected worldwide and a further 140 million individuals at risk of contracting the infection^1^. As a result of human migration and climate change, Chagas disease continues to expand beyond endemic zones to regions including North America, Europe and Australia, and it has been estimated that current levels of Chagas disease cause global economic hardship of $7.19 billion per year worldwide ^2^.

Most people with Chagas disease are asymptomatic and therefore unaware of the underlying high risk and remain untreated. Current treatment methods are effective for the acute infection but have toxic side-effects and are poorly effective against chronic infection^3^. Crucially, no vaccine nor robust and affordable diagnostic test is presently available for Chagas disease. Consequently, the World Health Organization has confirmed Chagas disease as a priority in its 2021-2030 Neglected Tropical Diseases roadmap^4^.

Current diagnostic tests have significant limitations including that they often detect infection after signs of the disease have already occurred in chronic infections, when current therapies are no longer effective. Some tests are also limited by shelf-life and require cold storage, resulting in diminishing sensitivity^5^. Additional limitations include cost, in the case of peptide-based tests, a lack of broad strain specificity, and a requirement for specialised personnel in resource-constrained health care settings (e.g., Elecsys ® Chagas test)^6 7 8^. The latter is often logistically difficult to implement in rural areas and is not affordable for some national healthcare systems^4^.

Thus, a practical and affordable solution is needed to address the health “ticking time bomb” that is Chagas disease. A new diagnostic test that can be easily employed in low-middle income countries, where cold chain storage of diagnostic reagents is not practical and expensive, would be of particular benefit.

Additionally, the development of rapid and affordable at-home or in-clinic diagnostic tests for Chagas disease would enhance privacy for receiving results, enabling individuals to detect the disease discreetly, given the stigma associated with the disease, who could seek timely medical intervention accordingly.

The *T. cruzi* Tc24 protein has been previously identified as a leading vaccine candidate for the development of Chagas disease^9^. Tc24 is a 24 kDa surface antigen^10^, also called FCaBP, for flagellum calcium binding protein, and is highly conserved in all genotypes of *T. cruzi* which cause Chagas disease^11^. With a sequence identity of more than 98% across strains^12^, the Tc24 polypeptide is an ideal target for the development of a broad-spectrum diagnostic test, overcoming the current limitations of genotype- and strain-specific tests. However, the wild-type (naturally occurring) Tc24 polypeptide has been previously reported to be prone to aggregation and showing limited expression in *E. coli* and *Pichia pastoris* expression systems^13^.

Mutagenesis of three cysteine residues to alanine residues in wild type Tc24 (3CS1 in NCBI Structure database but not fully characterised) has been previously performed in the attempt to reduce its limited stability and solubility. This led to stability up to 7 days at 4°C as reported in patent WO2017160849A1^14^. However, despite such improvement, a cold-chain would still be required and it did not solve the solubility and concentration problems to the extent needed to perform structural studies.

Therefore, here we addressed these problems by applying a computational approach, which led to highly thermostable and highly soluble Tc24 variants, which we tested *in vitro* using a plethora of biophysical techniques with an integrative approach. Strikingly, all variants also retained the ability of detecting antibodies in Chagas disease patient sera by ELISA confirming the retention of antigenicity. Additionally, transfer onto semi-solid lateral flow test (LFT) prototype confirmed antibody detection and therefore feasibility to develop a rapid diagnostic test.

## MATERIALS AND METHODS

### Protein design

Sequence conservation and phylogenetic analysis of Tc24 protein was performed based on the reference genome from CL Brener strain (NCBI gene reference XP_805575.1, Uniprot Q4CTX0) using BLASTP^15^. Sequence variation across *T. cruzi* strains was mapped onto the three-dimensional model of Tc24 generated with AlphaFold2 using CONSURF^16^ server. The partial structure of Tc24 has been determined before with PDB code 3CS1 from an unknown *T. cruzi* strain (Uniprot P07749), which is 99% identical to Miranda strain. Therefore, the corresponding PDB code of 3CS1 (in which only 17-208 aa are resolved), was used as experimental template for protein modification by PROSS2^17^. Three engineered variants were obtained from PROSS2, which were selected for further characterisation and named Tc24-V1, Tc24-V2 and Tc24-V3. The resulting PDB files from PROSS2 were extended at the N-terminal using AlphaFold2^18^ to add the missing N-ter and C-ter residue to complete the full 211 aa sequence based on the Miranda strain (Uniprot P07749), which has been reported as 3CS1_A in NCBI but without reference to the exact origin. PROSS2 design led to three variants ranked as the top heat-stabilised candidates: Tc24-V1 (K23S, S98A, A141T), Tc24-V2 (K23S, E61N, S72K, S98A, A141T, V156A, A170D, S194A) and Tc24-V3 (K23S, A44E, E61N, C66S, S72K, S98A, A141T, P157D, A161E, V166I, A170D, S194A, V196K).

### Protein expression and purification

The cDNA of Tc24 wild type (NCBI Structure entry 3CS1_A), CL Brener strain (Uniprot Q4CTX0) and PROSS2 engineered variants were all codon optimised for *E. coli* expression and sub-cloned into pET-15b expression vector with an additional N-terminal Spytag^19^ (TWIST). BL21(DE3) pLysS *E. coli* competent cells were transformed and selected on LB agar supplemented with 50 *µ*L/mL ampicillin. Cells were cultured overnight at 37°C in 2YT medium (Melford) supplemented with 100 *µ*g/mL ampicillin and inoculated 1:100 v/v in medium supplemented with the same antibiotic concentration and growth continued at 37°C. Once the culture reached an OD_600nm_ of 0.6, expression was induced by addition of IPTG to final concentration of 0.5 mM and continued for 21 hours at 20°C and under fixed parameters (30% Dissolved Oxygen, pH 7.2, 250rpm). Cells were pelleted by centrifugation at 4000xg for 15 min and resuspended in Buffer A (50 mM NaPi, 500 mM NaCl, pH 7.4) supplemented with 25 mM imidazole and EDTA-free protease inhibitor tablets (Roche). Cells were disrupted by sonication using a SoniPrep 150 Plus sonicator and soluble fraction obtained by centrifugation at 48,000xg for 30 min. Supernatant after centrifugation was loaded onto a Co^2+^-NTA resin (ThermoFisher), previously equilibrated with Buffer A and eluted in the same buffer supplemented with 500 mM imidazole. Protein expression was assessed by SDS-PAGE and Western blot, using a mouse anti-His6 HRP-conjugated antibodies (Invitrogen). Eluted protein samples were dialysed overnight against Buffer A and concentrated to 5 mg/mL using 10kDa MWCO concentrators (Millipore) before loading onto S200 16/600 (Cytiva) pre-equilibrated with 1x PBS. Protein-containing fractions were pooled together and concentrated to 10 mg/mL before being flash-cooled in liquid N2 for long-term storage at -80°C.

Expression media optimisation trials were performed by screening protein expression of Tc24 constructs in *E. coli* using: LB Broth (Miller), Glucose M9Y™ Hyper Broth™, Power Broth™, Superior Broth™ media from a Complete Media Optimization Kit™ (AthenaES, USA), along with 2x YT and Terrific Broth™ (Melford). Bacterial cultures were grown at 37°C in the respective medium supplemented with 50 µL/mL ampicillin and inoculated at 1:100 v/v in 100 mL shake-flasks of the same medium supplemented with the same antibiotic concentration. Growth was carried out at 37°C, 150rpm and protein expression induced when OD_600nm_ reached 0.6 by adding IPTG to a final concentration of 0.5 mM. Growth was then carried over for 21hrs at 20°C. Bacterial cultures were pelleted by centrifugation at 4000xg for 15 mins. The pellet was resuspended in 5 mL of buffer A per gram of bacterial pellet and lysis performed with a SoniPrep 150 Plus. Soluble fraction was separated from insoluble by centrifugation at 48,000xg for 30 min, and the remaining processing steps were performed as explained above.

Bioreactor expression of Tc24 engineered variants was performed in 10 L of Hyper Broth™(Molecular Dimensions, UK) supplemented with 50 µg/mL ampicillin and 50 µg/mL Antifoam 204 (Sigma) in an ElectroLab SIP 10L Vessel with ElectroLab FerMac 360 Tower, or AFS003 Bacterial System 15L TV vessel (InforsHT) fitted with EasyFerm Plus PHI K8 120 (pH control) (Hamilton), Growth was performed with constant parameters of 30% DO, pH 7.2, 250rpm, atmospheric pressure. Oxygen levels were maintained at an aeration rate of 1VVM and pH controlled with 1 M HCl (Sigma) and 1 M NaOH (Fisher). Bacterial culture was pelleted as described above, and frozen before further protein purification.

### Protein lyophilisation

Tc24 proteins were lyophilised overnight in 1x PBS (pH 7.4) at concentrations of ∼2 mg/mL, using a Labconco 710201050 freeze dryer (Labconco corp., USA) set at –85°C, 0.133mBar vacuum, for 18hrs. Samples were stored lyophilised at –80°C, and resuspended using equivalent volumes of sterile Milli Q H_2_O for further analysis.

### Biophysical characterisation

#### Size exclusion chromatography coupled with multiangle light scattering (SEC-MALS)

SEC-MALS experiments were performed using a Superdex 200 10/300 Increase column (Cytiva) and an AktaPure 25 System (Cytiva). The eluting protein was monitored using a DAWN HELEOS-II 18-angle light scattering detector (Wyatt Technologies), a U9-M UV/Vis detector (Cytiva), and an Optilab T-rEX refractive index monitor (Wyatt Technologies). Data were analysed using Astra v7 (Wyatt Technologies) with a refractive increment value of 0.185 mL/g.

#### Differential scanning fluorimetry using the inherent fluorescence of proteins (nDSF)

Temperature unfolding of Tc24-WT_3cs1_ Tag-less, Tc24-WT_3cs1_, Tc24-V1, Tc24-V2 and Tc24-V3 in PBS (pH 7.4) at a final concentration range of 80-100 μM, was performed in triplicate in Standard Prometheus NT.48 Capillaries, using the Prometheus NT.48 apparatus (NanoTemper, Munich, Germany) with the following parameters: excitation power: 70%, initial temperature: 20°C, final temperature: 95°C and temperature slope: 1.0°C/min.

### Circular Dichroism Analysis

All protein variants were prepared to a concentration ranging from 0.46-0.55 mg/ml. Comparative far UV CD experiments were performed using a nitrogen-flushed Module A end-station spectrophotometer with high throughput CD (HTCD) sample handling unit with 3 strings each containing 12 fused quartz 0.02cm pathlength cuvette cells, 1s integration time, 300mA ring current, 22°C at B23 Synchrotron Radiation CD Beamline at the Diamond Light Source, Oxfordshire, UK ^19,20^. The screening with HTCD enables the use of about 13 mL of solution per cell reducing substantially the volume of solution required compared to that using standard cuvelle cells. Overall the time required to measure the 36 samples and clean the strings was about 2.5hrs without the need for repeated cleaning between samples for single cell measurements ^21-24^. Temperature studies were carried out on Chirascan Plus CD Spectropolarimeter (Applied Photophysics Ltd). All samples were incubated at 20°C and spectra measured every 5°C steps in a temperature range between 20° and 90°C with 2 min equilibration time for each temperature gradient. Reversibility was monitored by measuring the spectrum at 20°C after cooling from 90°C with 10 minutes incubation time. Data analysis was performed using CDApps ^25^. Additional spectra were recorded for the variants at fixed temperature of 22°C in the following buffers: 10 mM HEPES, 75 mM NaCl (pH 7.4); 10 mM MOPS, 75 mM NaCl (pH 7.4); 10 mM Glycine, 75 mM NaCl (pH 3.6); 50 mM NaPi (pH 7.4); 20 mM Sodium Acetate (pH 5.6); 10 mM Tris, 1 mM EDTA (pH 7.4); 0.5X PBS (pH 7.4); and 0.5X TBS (pH 7.4).

### 1D ^1^H -NMR

1D ^1^H-NMR spectra of Tc24 proteins at concentrations of ∼100 μM were recorded in PBS buffer. Protein stability in solution was monitored over time, at regular time intervals using successive ^1^H one-dimensional (1D) spectra. The 1D ^1^H spectra were recorded for a period of 70 days at 303K, whilst storage was performed at 278K.

The time between the addition of the complex and the acquisition of the first 1D ^1^H spectrum was monitored (68 s to 81 s, depending on the peptide) and noted down for each experiment. NMR spectra were acquired using a Bruker Avance Neo 800 MHz spectrometer with a 5mm, z-gradient TXO cryoprobe, and were processed using TopSpin (Bruker). The integrals of the peaks of interest were determined using the Kinetics Method in the program Dynamics Center (Bruker).

### Flow Induced Dispersion Analysis (FIDA) experiments

FIDA measurements were taken using a FIDA 1 instrument equipped with a Fidabio 280 nm LED fluorescence detector (Fida Biosystems ApS). A standard silica capillary (i.d.: 75 μm, LT: 100 cm, Leff: 84 cm) coated with HS reagent (Fida Biosystems ApS) was used for all experiments. The experiments were performed in 1x PBS (pH 7.4) at 25°C and the viscosity of the 1x PBS (0.00089 Pa*s) was determined experimentally using a Lovis 2000 ME microviscometer (Anton Paar). The following stepwise procedure was applied for the FIDA experiments. Milli Q H2O was used for flushing the HS-coated capillary at 3500 mbar for 60 s. The capillary was then equilibrated using 1x PBS at 3500 mbar for 40 s followed by injection of the purified protein at 50 mbar for 10 s. Lastly, the purified protein was mobilized to the fluorescence detector with 1x PBS at 400 mbar for 240 s. The equilibrium dispersion profile (Taylorgram) was recorded in the latter step. The Taylorgrams were analysed, with viscosity compensation, using FIDA software (V3.0.2, Fida Biosystems ApS) to determine the apparent hydrodynamic radius for each condition run. Purified protein samples run were at a concentration range of 80-100 μM, in triplicate. For freeze-thaw cycle preparation, protein aliquots were thawed for 1 min at 42°C in a heated water bath, flash-frozen in Liquid N2, and either stored at -80°C or repeated 5 times.

### ELISA experiments

No new human samples were collected specifically for this work. All human sera used here are previously archived, with consent for research, anonymised, coded, and do not reveal patient identities^20,21^ Immulon 4HBX plates (735–0465, VWR) were coated with 100 µL/well of recombinant Tc24 protein at 1 µg/mL in coating buffer (15 mM Na_2_CO_3_, 34 mM NaHCO3, pH 9.6). After overnight incubation at 4°C, plates were washed three times with PBS/0.05% (v/v) Tween 20 (PBST), then blocked with 200 μL of PBS/2% skimmed milk powder at 37°C for 2 h. Following three washes, 100 μL of human serum samples diluted 1:200 v/v in blocking buffer were added and incubated for 1 h at 37 °C. After six washes, 100 μL/well of horseradish peroxidase (HRP)-conjugated donkey anti-human IgG (H+L) antibody (709–035–149, Jackson Immuno Research) diluted 1:2,000 v/v in blocking buffer was added and incubated for 1 h at 37 °C. After six washes, plates were developed using 100 μL/well of 2 mM o-phenylenediamine dihydrochloride (OPD) in phosphate-citrate buffer (pH 5.0) containing 0.005% H_2_O_2_. Reactions were stopped after 10 min by addition of 50 μL/well of 2 M H_2_SO_4_, and absorbance was measured at 490 nm. using a plate reader (Dynatech MRX). Experiments were conducted in duplicate in separate ELISA plates.

### Prototype Lateral Flow test (LFT)

Lateral-flow tests were manufactured with overlapping absorbent upper pad, nitrocellulose membrane, conjugate and sample pads, all backed with a plastic strip, as previously described^21^. The nitrocellulose membrane (CN95, Sartorius) was sensitized with biotinylated Tc24 antigen coupled to avidin (test line, T), goat antibody reagent (control line, C) and reduced Triton X100 (Sigma-Aldrich, deposit line, D). Detection conjugate (Protein G coupled to colloidal gold particles) was dried on fiber glass conjugate pad. Strip is inserted in a plastic cassette. The cassette has two windows: buffer well and test/reading window.

Depending on sample type, 2 (serum/plasma)-3 (whole blood) µl of sample was applied the deposit line (D), immediately followed by 5µl of running buffer on the same line. Then 120 µl of running buffer was added in the buffer well. IgG antibodies from the sample that migrated in the strip first encounter and react with the protein G gold conjugate rehydrated by the buffer and migrate further over the nitrocellulose membrane and reacted with the avidin-immobilized Tc24 antigen (T line), resulting in a red-purple colored band. The remaining conjugate further migrates and reacts with the immobilized reagent at the control line (C) ensuring that sample and conjugate migration had occurred. Tests were read after 15 minutes and scored as either positive (coloured test and control lines) or negative (coloured control line only) and photographed. Samples for testing cross-reactivity with visceral leishmaniasis and malaria are from Sudanese patients with malaria or visceral leishmaniasis, from LSHTM, collected as part of the NIDIAG project^22^.

## Results and discussion

Tc24 is a leading candidate for therapeutic and diagnostic solutions for Chagas disease^23^. Despite this, limited stability and protein expression scalability have hampered the field. The only Tc24 structure information, which is essential for computational design, belongs to an unknown strain of *T. cruzi*, which is 99% identical to Miranda strain (PDB code 3CS1). We, decided to clone and express Tc24 from the well-characterised and reference strain of *T. cruzi* CL Brener, namely Tc24-WT_Q4CTX0_. Tc24-WT_Q4CTX0_ was cloned, expressed and purified wild-type Tc24 as an additional control for our study (Fig.1).

**Fig. 1.**
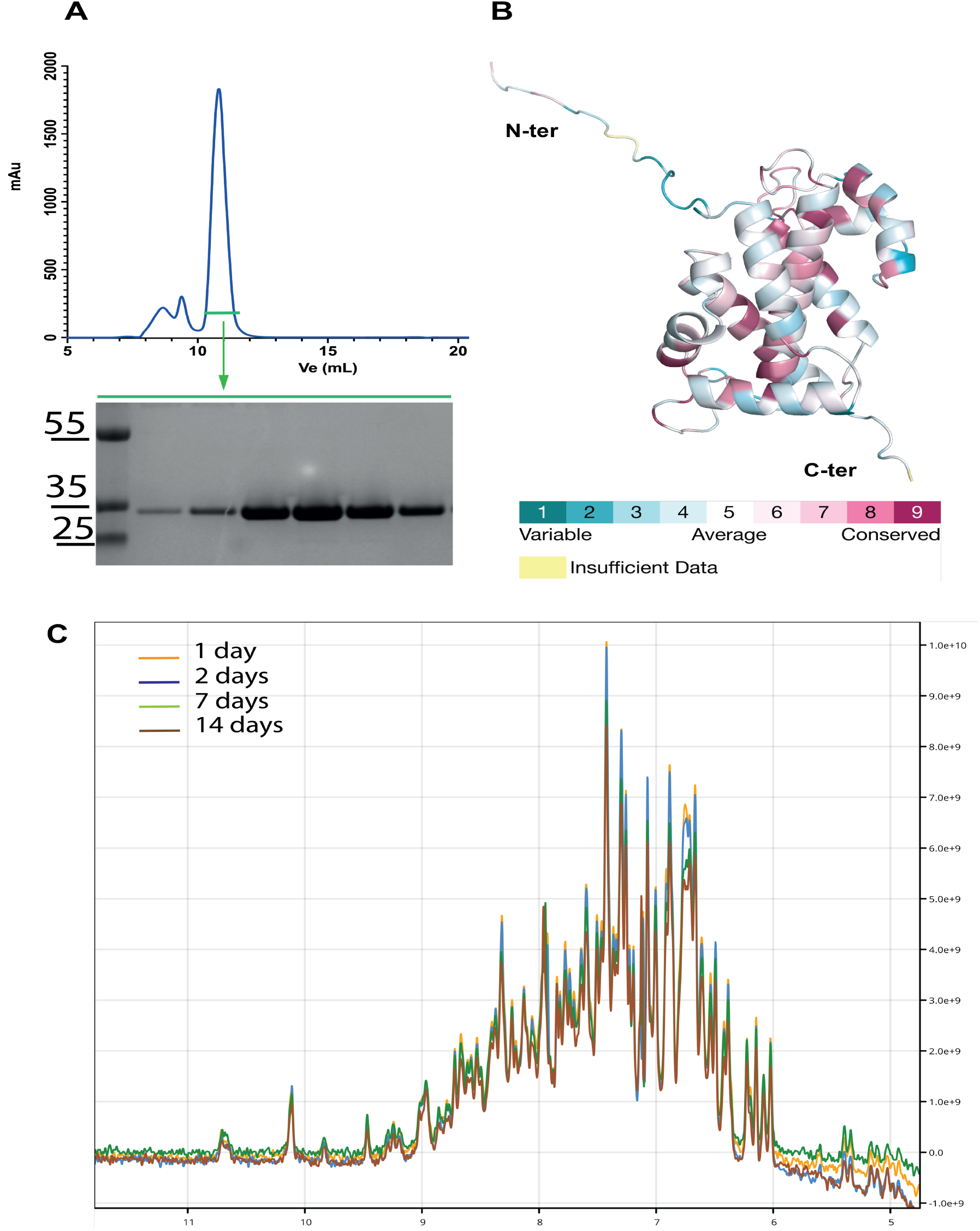
Wild-type Tc24 Spytagged and His tagged construct (Tc24-WT_Q4CTX0_) purification and in solution 1D-NMR analysis. (**A**) Recombinant Tc24 was purified by affinity chromatography followed by SEC with S200 10/300. (**B**) CONSURF model showing the sequence conservation of Tc24 mapped onto the full-length Tc24 structure obtained with AlphaFold2. Colour coding corresponds to sequence conservation. (**C**) Protein stability was recorded every 7 days, up to 14 days at 20°C.

We applied a computational approach to overcome existing limitations of Tc24, using the amino acid sequence of Tc24 (PDB ID:3CS1), as an input for PROSS2.

The top three engineered variants (V1, V2, V3), as ranked in silico by PROSS2, were selected and harbour between three and fourteen mutations, respectively: Tc24-V1 (K23S, S98A, A141T), Tc24-V2 (K23S, E61N, S72K, S98A, A141T, V156A, A170D, S194A) and Tc24-V3 (K23S, A44E, E61N, C66S, S72K, S98A, A141T, P157D, A161E, V166I, A170D, S194A, V196K) (Fig.2), which were located across all alpha helices except alpha helix 6 and mapped versus the wild-type templates (Fig.S1A-B). All variants were recombinantly expressed in *E. coli* (Fig.3) and purified by SEC. All the variants were eluted at expected MW, as further confirmed by SEC-MALS analysis (Fig. 4).

**Fig. 2.**
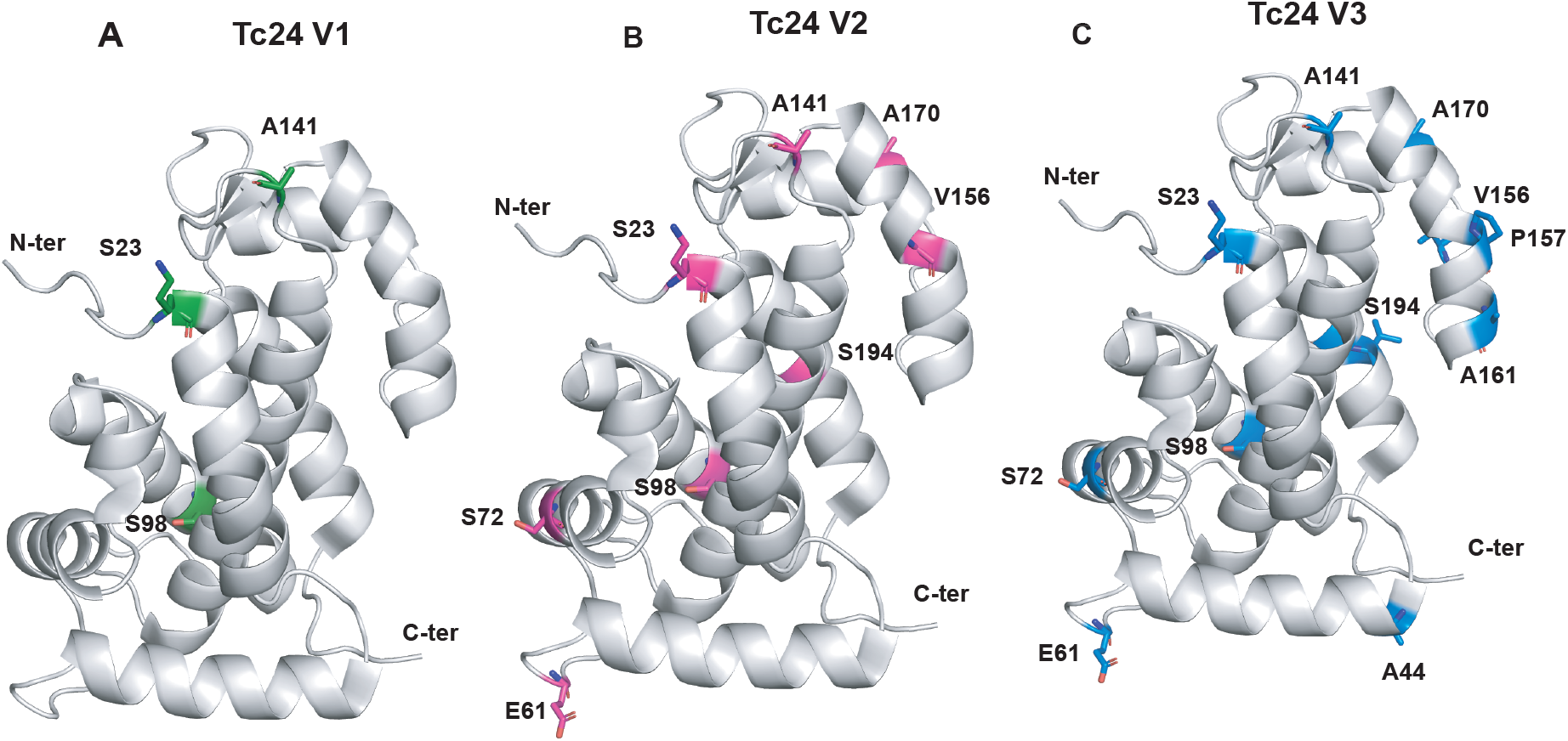
Mapping of mutated amino acid positions onto the crystal structure of wild-type Tc24. (PDB ID 3CS1). Amino acid positions which were mutated are mapped onto the wild-type structure of Tc24 (3CS1), Tc24-V1 (A), Tc24-V2 (B) and Tc24-V3 (C) and coloured in green, magenta and blue, respectively. The side-chains of the mutated amino acids are shown in sticks representation coloured by atom type using PyMOL 2.5.3.

**Fig.3.**
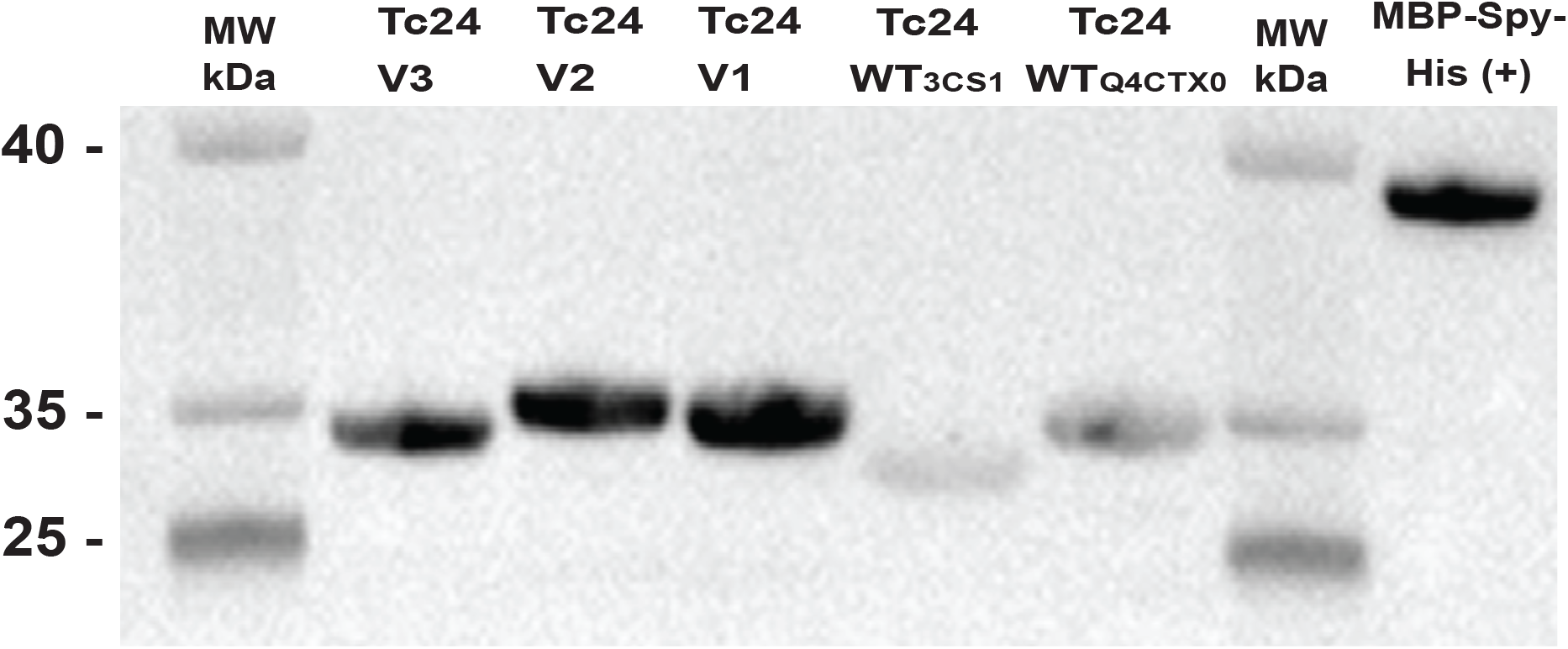
Protein expression of Tc24 variants. Western blot analysis of protein expression yields across Tc24 variants using anti-His antibody and comparison with Tc24-WT_3CS1_ and Tc24-WT_Q4CTX0_. MBP construct with His and Spy-tags was used as control.

**Fig.4.**
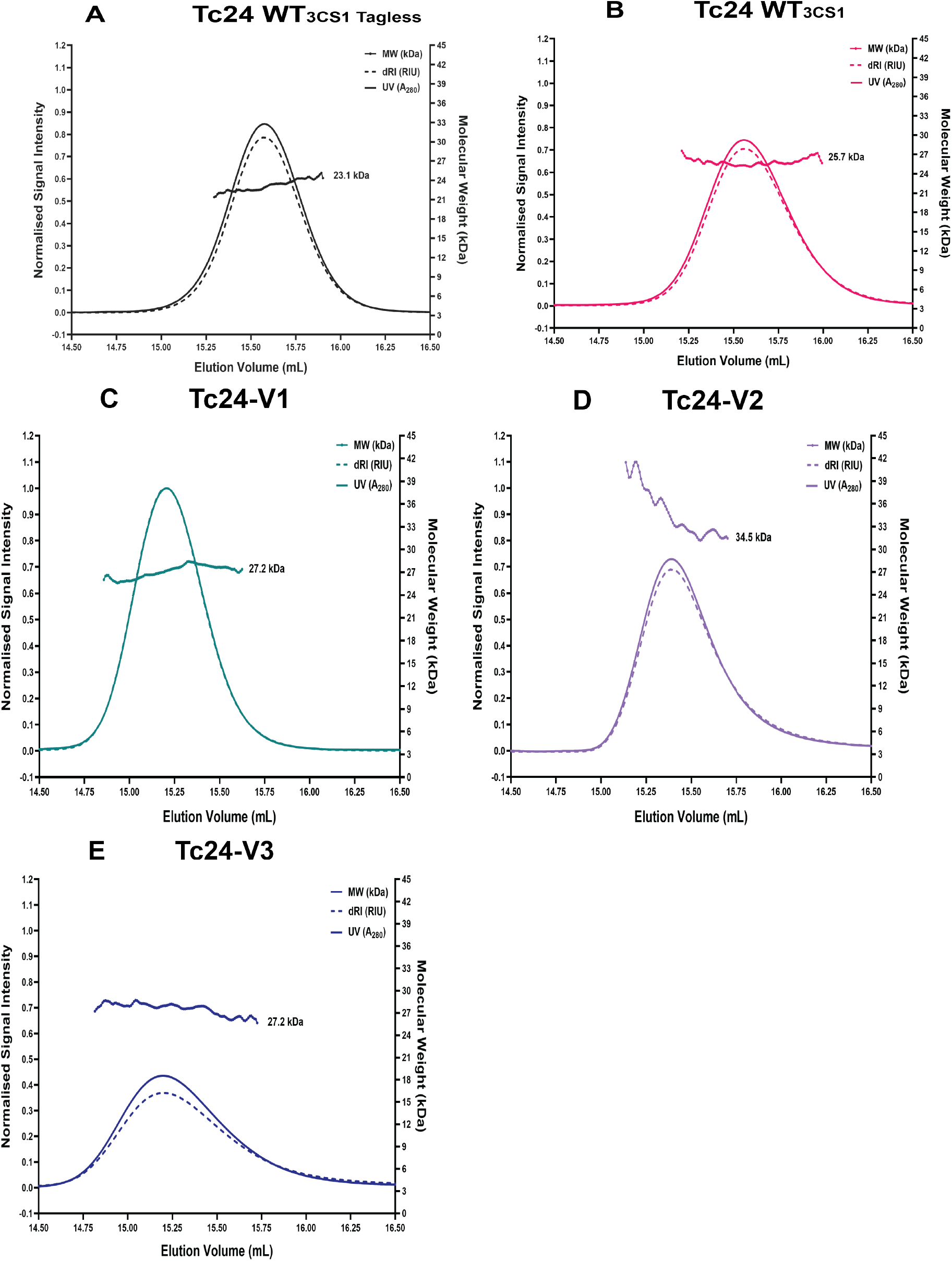
SEC-MALS analysis of Tc24 Variants. The UV A_280_ (UV) data and differential refractive index (dRI) data recorded for each of the Tc24 constructs is shown. The molecular weight (MW, kDa) distribution across each peak is also indicated and, in all cases, this is consistent with each construct behaving as a monomer in 1X PBS (pH 7.4) solution. Tc24-WT_3CS1_ Tag-less (A), Tc24-WT_3CS1_ (B) and variants Tc24-V1 (C, green), Tc24-V2 (D, violet), Tc24-V3 (E, blue).

The variants were initially produced in batch *E. coli*, to identify the best growth conditions (Fig.S2) before scaling-up the protein production in biofermenters with resulting protein yield exceeding 40 mg per litre of fermentation. Protein expression of the variants was optimised to produce up to five-fold more protein compared to the reported wild-type constructs (Fig.3), in which tags were also removed to produce tag-less variants (Fig.S3). To evaluate the improved properties of our three variants compared to the wild-type proteins from CL Brenner and 3CS1_A, respectively named Tc24-WT_Q4CTX0_ or Tc24-WT_3cs1_, we adopted a plethora of complementary biophysical techniques. First, we confirmed their monodispersion in solution using SEC-MALS (Fig.4) and then compared their hydrodynamic radii through Flow Induced Dispersion Analysis (FIDA) (Fig.5A). To test stability across repeated freeze-thaw cycles, we applied FIDA analysis, which showed no significant difference in variant hydrodynamic radius between freeze-thaw cycles 1 to 5 (Fig.5B). In addition, label-free thermal stability experiments showed a clear increase in melting temperature (T_m_), with the increase in thermal stability correlating with the increasing number of engineered amino acids in the Tc24-WT_3cs1_ construct: Tc24-V1 (T_m_ = 57.0 ± 1.46 °C), Tc24-V2 (T_m_ = 63.3 ± 0.21 °C), and Tc24-V3 (T_m_ = 73.6 ± 0.15°C). WT (T_m_ = 53.5 ± 0.12 °C) and tag-less WT (T_m_ = 54.6 ± 0.31 °C) Tm values were comparable, indicating intrinsic heat-tolerance is purely sequence modification driven (Fig.6) CD-melt experiments allowed assessment of the heat-stability in solution of each variant, measured as the loss of secondary structure elements, showed an unfolding temperature of 60.4, 67.4 and 69.2°C for variants Tc24-V1, Tc24-V2 and Tc24-V3, respectively (Fig.7).

**Fig. 5.**
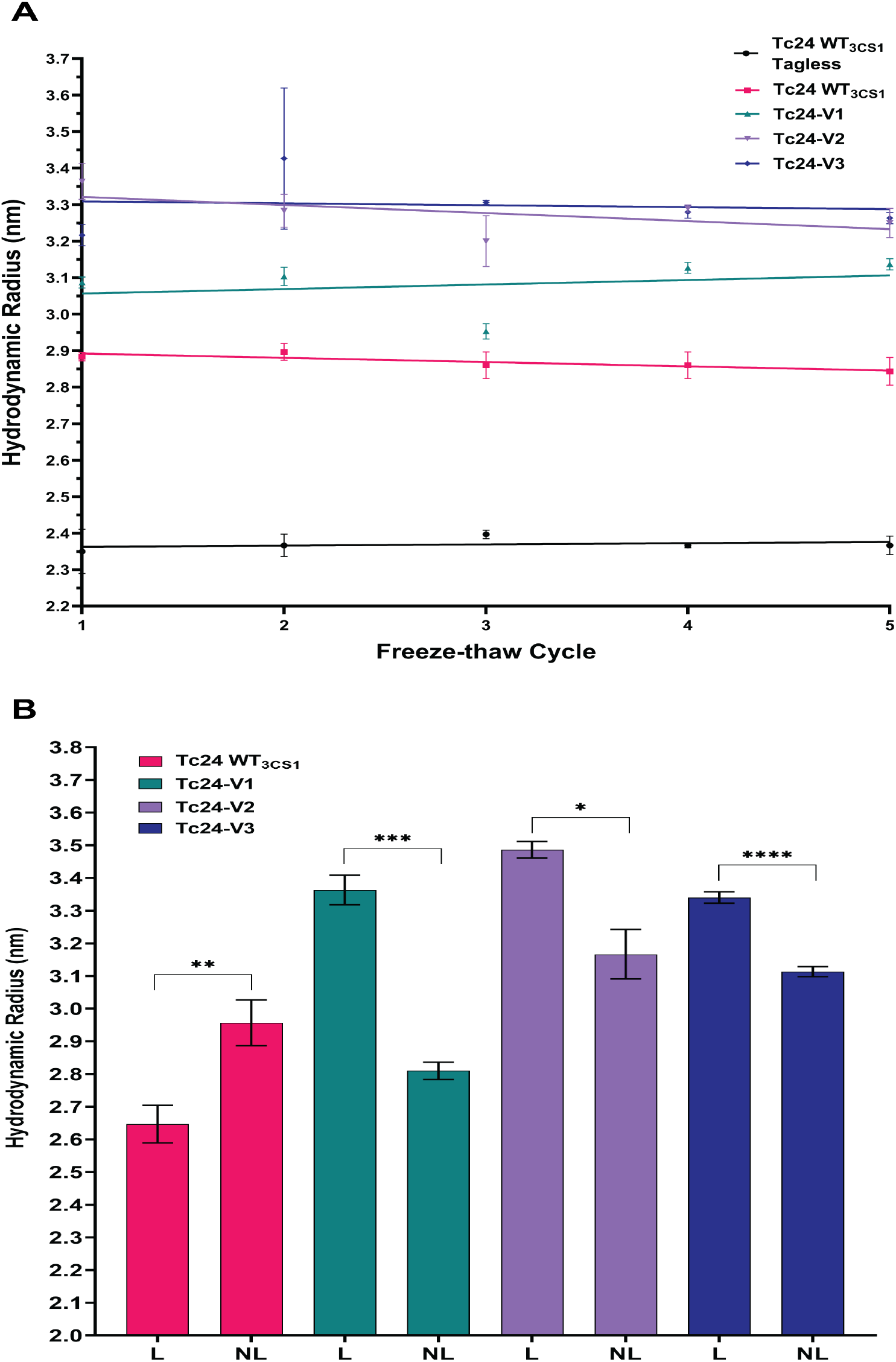
FIDA analysis of Tc24 variants. **A**. Hydrodynamic radius (nm) changes were recorded across multiple flash-cooling thawing cycles with liquid N2 for Tc24 WT, Tc24 WT His-tagged and Tc24 variants TC24-V1 (green), TC24-V2 (light violet) and TC24-V3 (dark violet). A two-way RM ANOVA revealed no significant change of Rh by freeze-thaw cycle (F(2.441, 34.17) = 2.26, P = 0.11; Geisser-Greenhouse corrected). **B**. Hydrodynamic radius (nm) changes were recorded before (NL) and after lyophilisation (L) conditions. Data are presented as mean ± SEM. Statistical analysis was performed using Welsh’sT-test (Tc24 WT His tagged t(3.856) = 5.906, P = 0.0046, two-tailed; Tc24-V1 t(3.231) = 18.33, P = 0.0002; Tc24-V2 t(2.437) = 6.946, P = 0.0116; Tc24-V3 t(2.018) = 2.488, P = 0.0087). All experiments were performed in 1X PBS pH 7.4.

**Fig. 6.**
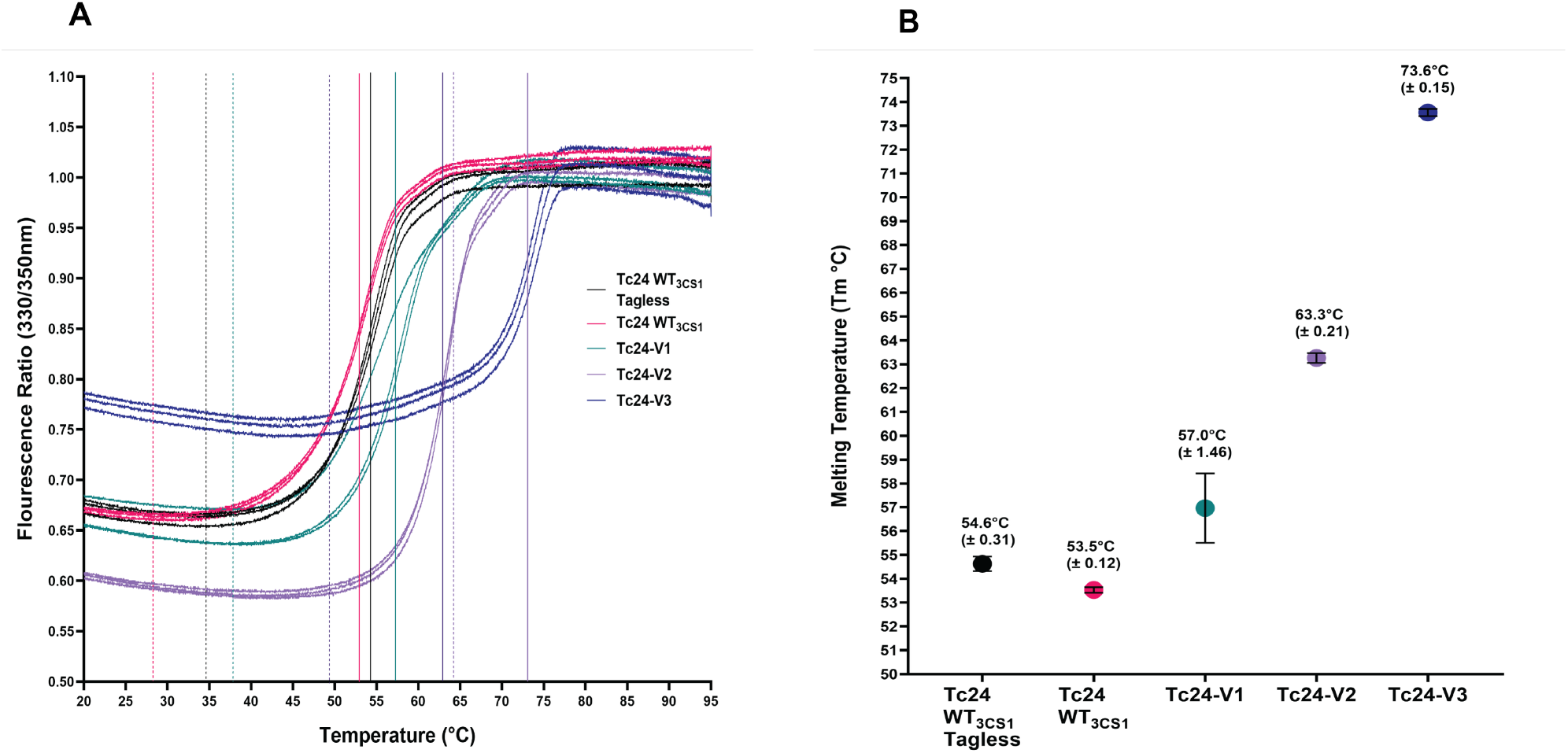
Heat stability of Tc24 variants by nanoDSF. A. The thermograms recorded for each Tc24 construct are shown together with lines indicating the unfolding onset points (dotted vertical lines) and melting temperatures (T_m_) values (solid vertical lines). B. T_m_ values were recorded for all variants as function of the intrinsic tryptophan fluorescence ratio (330/350 nm) for Tc24-WT_3CS1_ Tag-less (black), Tc24-WT_3CS1_ (red), and variants Tc24-V1 (green), Tc24-V2(violet) and Tc24-V3 (blue). Experiments were performed in 1X PBS (pH 7.4).

**Fig. 7.**
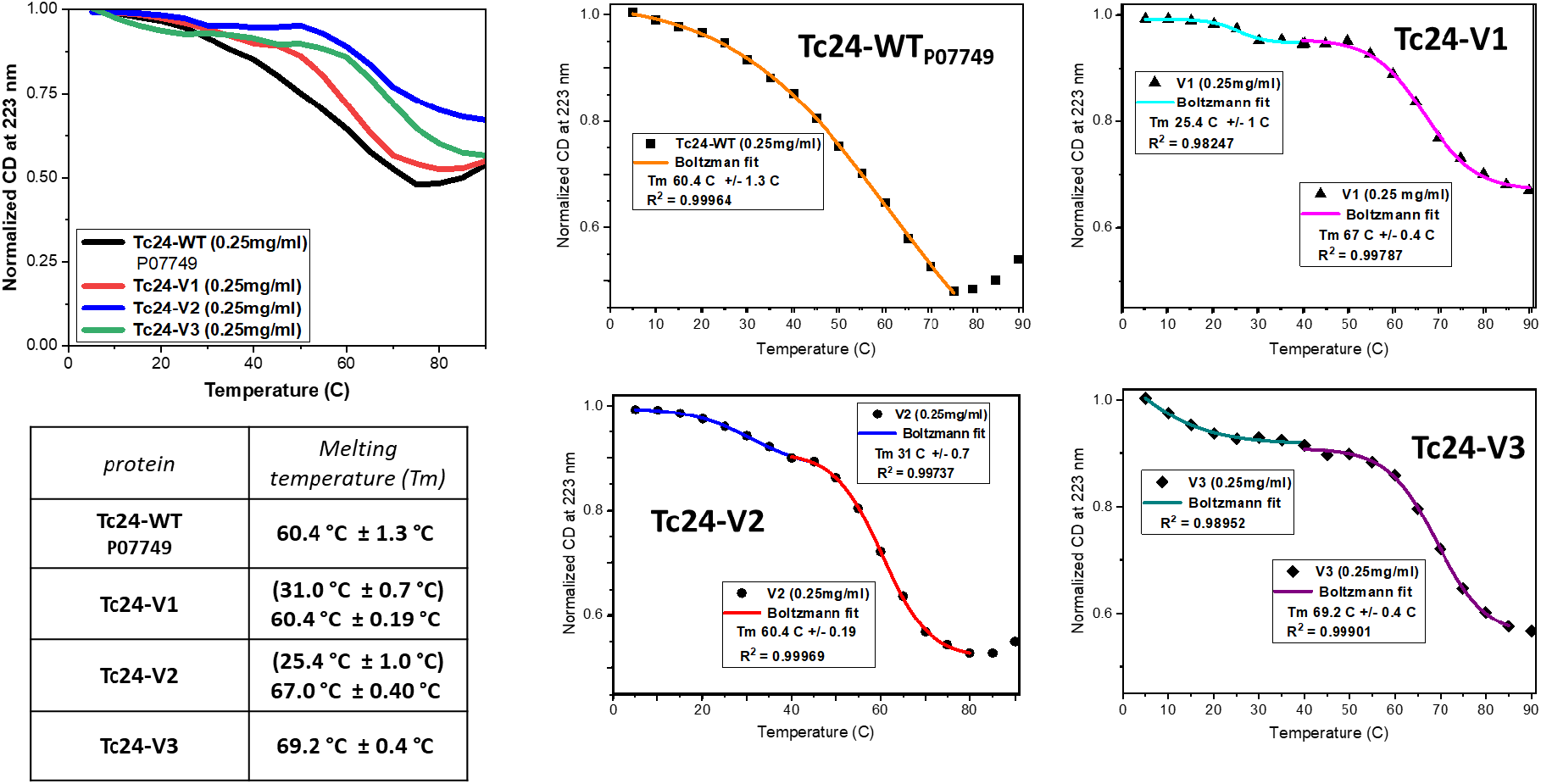
CD-melting measurements of Tc24 WT and Tc24 variant heat-stability. CD spectra were recorded as a function of temperature in the far UV region. Change in CD intensity at 223 nm were normalised and plotted to calculate the T_m_ with OriginPro^®^ using the Boltzmann equation^27,28^.

Additionally, the variants were stable across a range of standard purification buffers with the most stable variant appearing to be Tc24-V2 (Fig. S4).

Strikingly, 1D ^1^H -NMR experiments showed an astonishing stability of the recombinant Tc24 proteins in solution to up to 70 days at 25°C (Fig.8). The Tc24-WT3cs1 tagged protein started unfolding after 12 days, whilst all variants remained stable in solution for the same amount of time as indicated by the disappearance or broadening of the typical amide group peak at around 7 ppm over time.

**Fig. 8.**
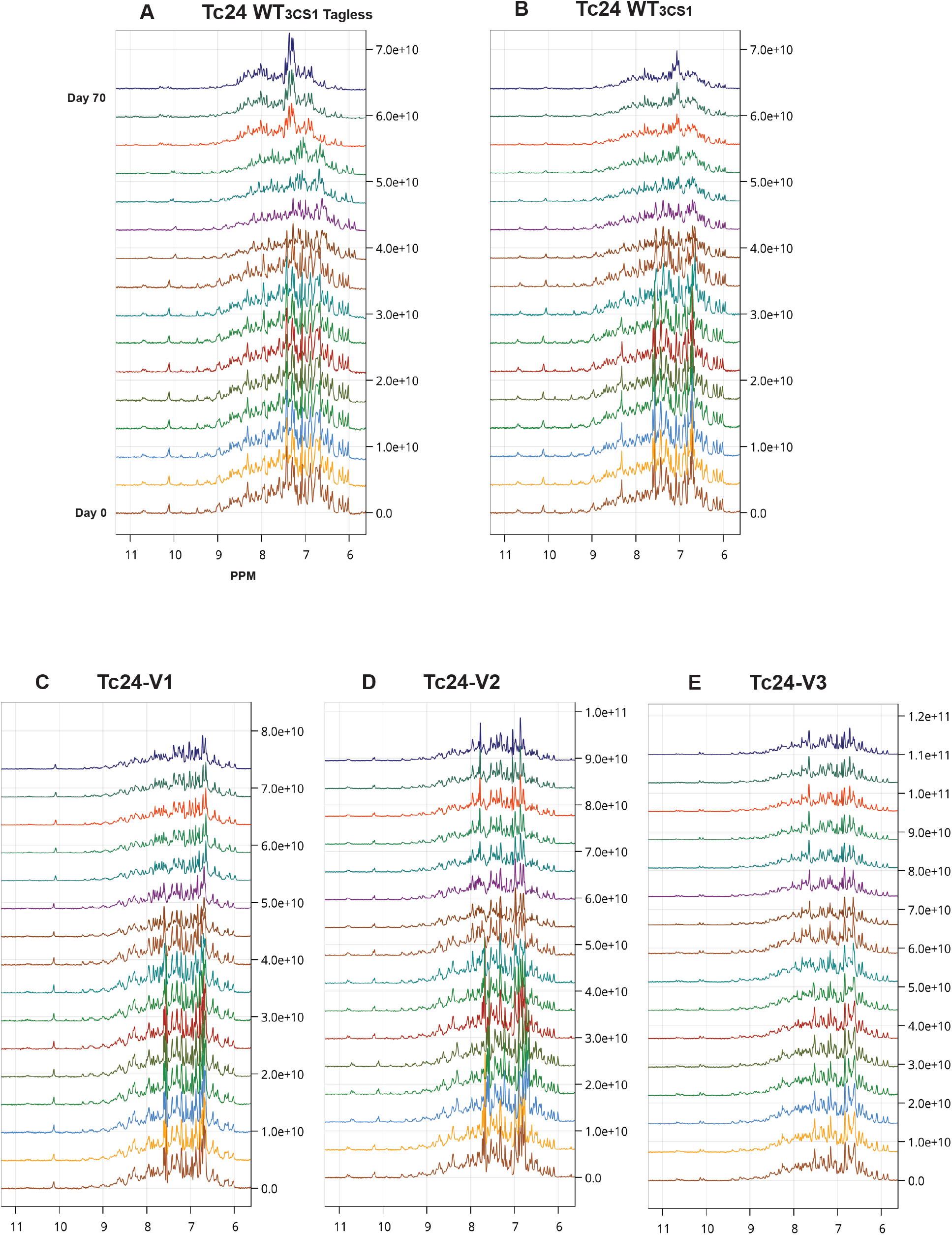
^1^H-NMR analysis of stability in solution over time of Tc24 variants. ^1^H-NMR experiments recorded on an 800 MHz spectrometer at 303K at different time points in PBS buffer show stability up to 69 days. NMR experiments were performed at the Leicester Institute of Structural and Chemical Biology (LISCB, University of Leicester). Protein storage was at 278K for the entire length of the experiment.

Additionally, lyophilisation experiments also confirmed that all the variants their folding (Fig. 5B and Fig. S5), as well as showed by the evaluation of their monodispersity in-solution with comparable hydrodynamic radii and molecular weight ranges.

ELISA experiments were conducted to compare the performance of the Tc24 variants versus wild-type Tc24 from CL Brener strain construct in detecting naturally occurring antibodies in Chagas disease patients from the anonymised LSHTM Biobank versus healthy donors. Our results show that the antigenicity of the improved variants remain unaltered, as similar performance to wild-type Tc24 was established (Fig.9). As expected, the strength of response across samples from patients from the southern regions of Latin America produced higher signal in ELISA and LFT compared to those from (Fig.9) the North Cone. This was not unexpected as the antibody titres remain higher in the South Cone where the disease is endemic^24^.

**Fig. 9.**
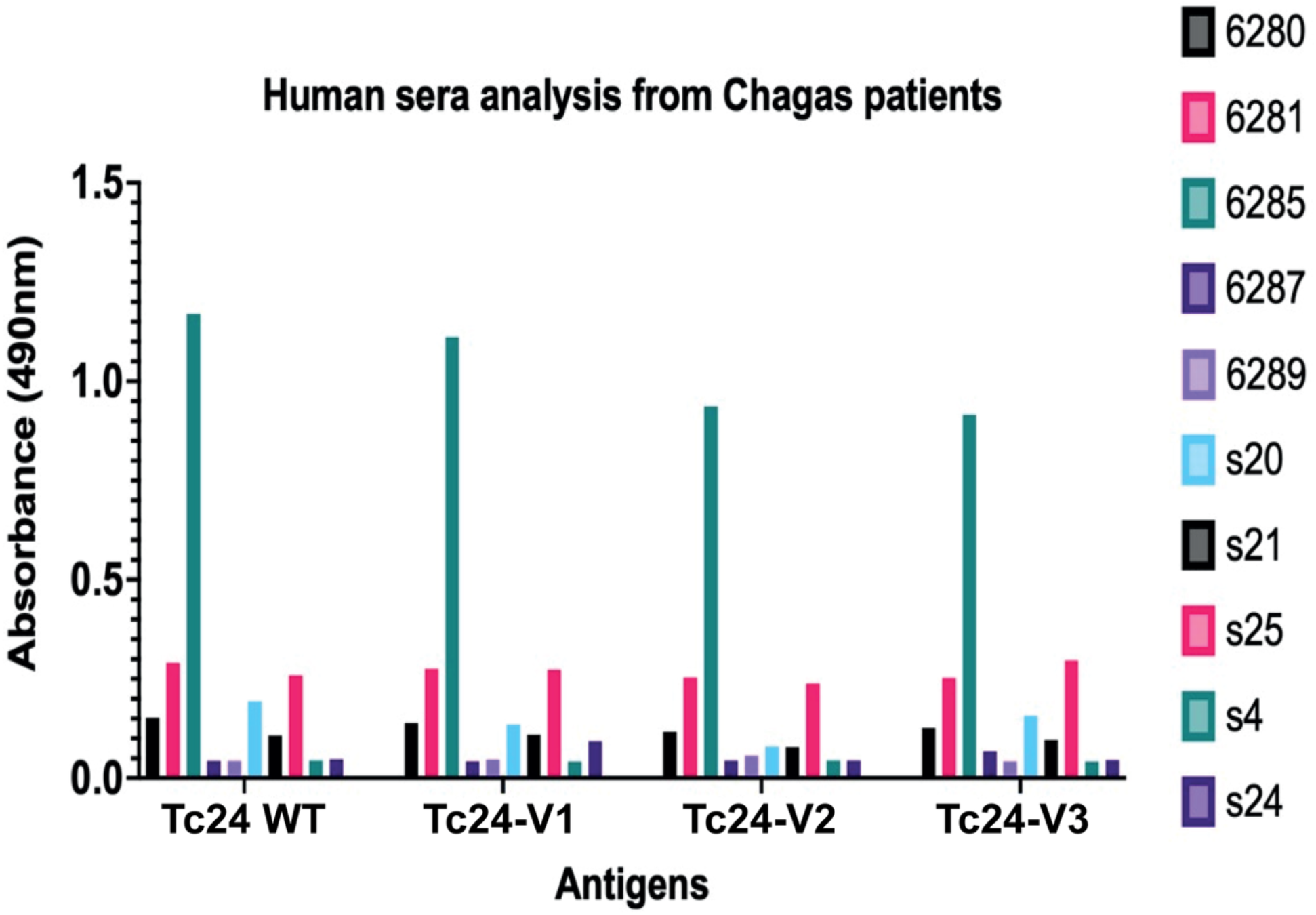
ELISA testing against human sera from Chagas patients from Northern and Southern cones of Latin America. All antigens were screened against Chagas patients’ sera from Northern (“6” codes) and Southern (“S” codes) hemispheres from LSHTM Biobank by ELISA using antigen at 0.7 micrograms/mL (B). Data were normalised compared to healthy donors. Samples identifiers are anonymised with numbers and/or letters.

To assess feasibility and the retention of antibody detection from patients when transitioning from liquid to semi-solid state, we immobilised Avi-tagged Tc243cs1 onto nitrocellulose strips enclosed in LFT cassettes and applied the samples from the highest titre from the southern region (S1 and S2). In parallel, we assessed in duplicate the specificity to detect Chagas disease versus malaria or leishmaniasis. Our results showed that no false positive were detected with this approach (Fig10.)

**Fig.10.**
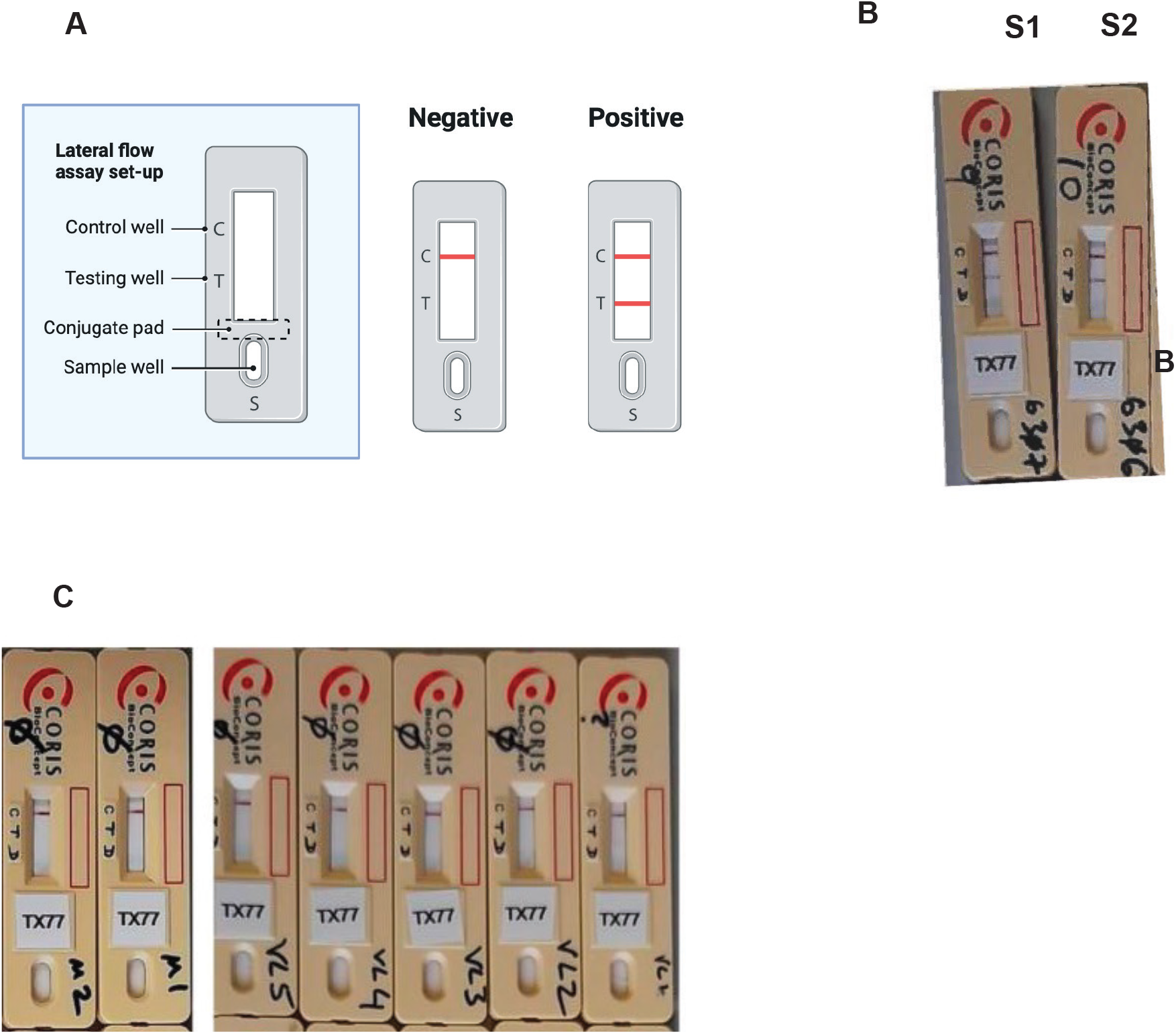
Preliminary feasibility study for transfer to semi-solid state of wild-type Tc24. Schematic Lateral Flow Test (LFT) model (Biorender license https://BioRender.com/y9gxqwm) **(a)**. Test of patient sera from South America (S1, S2) on the LFT prototype **(b)**. Wild-type Avi-tagged Tc24 (named TX77) was transferred to nitrocellulose strips and placed in LFT cassettes. Cross-reactivity with samples from patients with malaria and leishmaniasis was also performed indicating no false positives. Samples are from Sudanese patients with malaria or visceral leishmaniasis, from LSHTM, collected as part of the NIDIAG project^22^ **(c)**.

Altogether, our results demonstrate the feasibility of transferring the most heat-tolerant and most stable variant over time (Tc24-V2) to a membrane bound antigen specific test for further testing on a Lateral Flow Test (LFT) (patent pending PCT 2416406.3) across a larger number of samples from the Northern and Southern regions of Latin America.

## Author Contributions Statement

S.B. and C.W. expressed and purified proteins, characterised them biophysically. S.B., C.W. prepared the samples for NMR acquisition and F.M. collected spectra and analysed them. G.H. collected with C.W. nDSF, SEC-MALS and FIDABIO data and analysed them. G.S., R.H., T-M.G. collected and analysed CD data. T.B. generated and analysed ELISA and LFT data. L.T., M.D., P.M. and B.R. provided advice and guidance for ELISA and LFT optimisation. I.C. conceptualised the experiments and provided funding. S.B., C.W. and I.C. prepared the manuscript and all authors contributed and commented on it.

## Acknowledgements

We would like to acknowledge staff at B21 at the Diamond Light Source for support for data collection, B23 beamline access project number SM40707. The project was funded by: Wellcome grant 204801/Z/16/Z (IC) and by MRC IAA institutional grant to the University of Nottingham (MR/X502741/1).

