## Supplementary Data for "Engineered heat-stable variants of *Trypanosoma cruzi* flagellar protein Tc24 enable serological detection of Chagas disease"

### Supplementary Information

Tc24 (3CS1) Topology

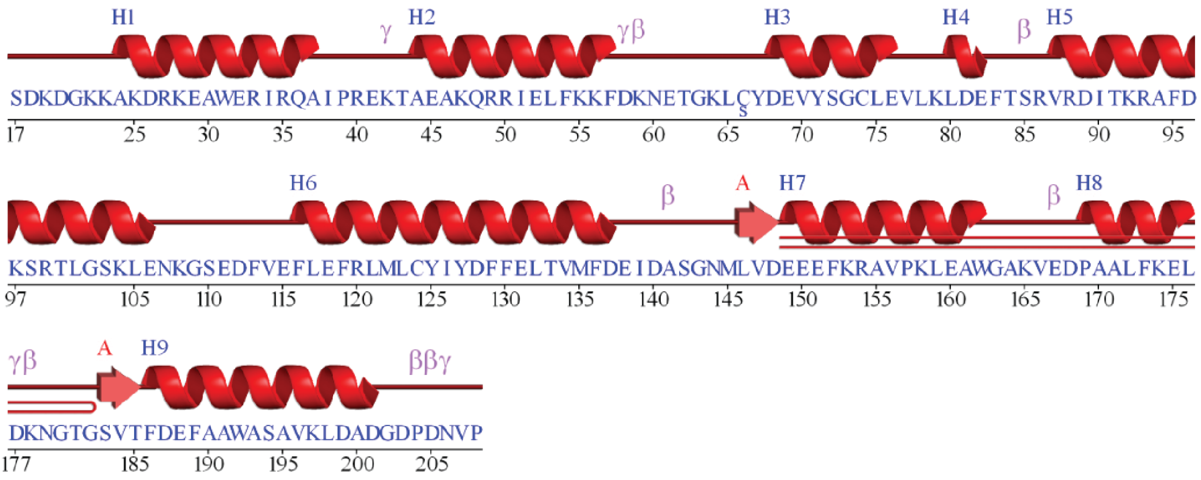


```
                   1                                                     55
Tc24-WT  Q4CTX0  MGACGSKDSTSDKGLASDKDGKNAKDRKEAWERIRQAIPREKTAEAKQRRIELFK
Tc24-WT  P07749  MGACGSKGSTSDKGLASDKDGKKAKDRKEAWERIRQAIPREKTAEAKQRRIELFK
Tc24-V1          MGACGSKGSTSDKGLASDKDGKSAKDRKEAWERIRQAIPREKTAEAKQRRIELFK
Tc24-V2          MGACGSKGSTSDKGLASDKDGKSAKDRKEAWERIRQAIPREKTAEAKQRRIELFK
Tc24-V3          MGACGSKGSTSDKGLASDKDGKSAKDRKEAWERIRQAIPREKTEEAKQRRIELFK

                   56                                                   110
Tc24-WT  Q4CTX0  KFDKNETGKLCYDEVHSGCLEVLKLDEFTPRVRDITKRAFDKARALGSKLENKGS
Tc24-WT  P07749  KFDKNETGKLCYDEVHSGCLEVLKLDEFTPRVRDITKRAFDKARALGSKLENKGS
Tc24-V1          KFDKNETGKLCYDEVYSGCLEVLKLDEFTSRVRDITKRAFDKARTLGSKLENKGS
Tc24-V2          KFDKNNTGKLCYDEVYKGCLEVLKLDEFTSRVRDITKRAFDKARTLGSKLENKGS
Tc24-V3          KFDKNNTGKLSYDEVYKGCLEVLKLDEFTSRVRDITKRAFDKARTLGSKLENKGS

                   111                                                  165
Tc24-WT  Q4CTX0  EDFVEFLEFRLMLCYIYDFFELTVMFDEIDASGNMLVDEEEFKRAVPKLEAWGAK
Tc24-WT  P07749  EDFVEFLEFRLMLCYIYDFFELTVMFDEIDASGNMLVDEEELKRAVPKLEAWGAK
Tc24-V1          EDFVEFLEFRLMLCYIYDFFELTVMFDEIDTSGNMLVDEEEFKRAVPKLEAWGAK
Tc24-V2          EDFVEFLEFRLMLCYIYDFFELTVMFDEIDTSGNMLVDEEEFKRAAPKLEAWGAK
Tc24-V3          EDFVEFLEFRLMLCYIYDFFELTVMFDEIDTSGNMLVDEEEFKRAVDKLEEWGAK

                   166                                     211
Tc24-WT  Q4CTX0  VEDPAALFKELDKNGTGSVTFDEFAAWASAVKLDADGDPDNVPESA
Tc24-WT  P07749  VEDPAALFKELDKNGTGSVTFDEFAAWASAVKLDADGDPDNVPESA
Tc24-V1          VEDPAALFKELDKNGTGSVTFDEFAAWASAVKLDADGDPDNVPESA
Tc24-V2          VEDPDALFKELDKNGTGSVTFDEFAAWAAAVKLDADGDPDNVPESA
Tc24-V3          IEDPDALFKELDKNGTGSVTFDEFAAWAAAKKLDADGDPDNVPESA
```

**Fig. S1 Protein topology for Tc24 3CS1** with helix boundary position delimitation, generated by PDBsum web server (top). Multiple sequence alignment of Tc24 $WT_{P07749}$ and Tc24 $WT_{Q4CTX0}$ against V1-3 highlighting PROSS2 amino acid substitutions in V1 (Green), V2 (Green, purple), V3 (Green, purple, blue) (bottom).

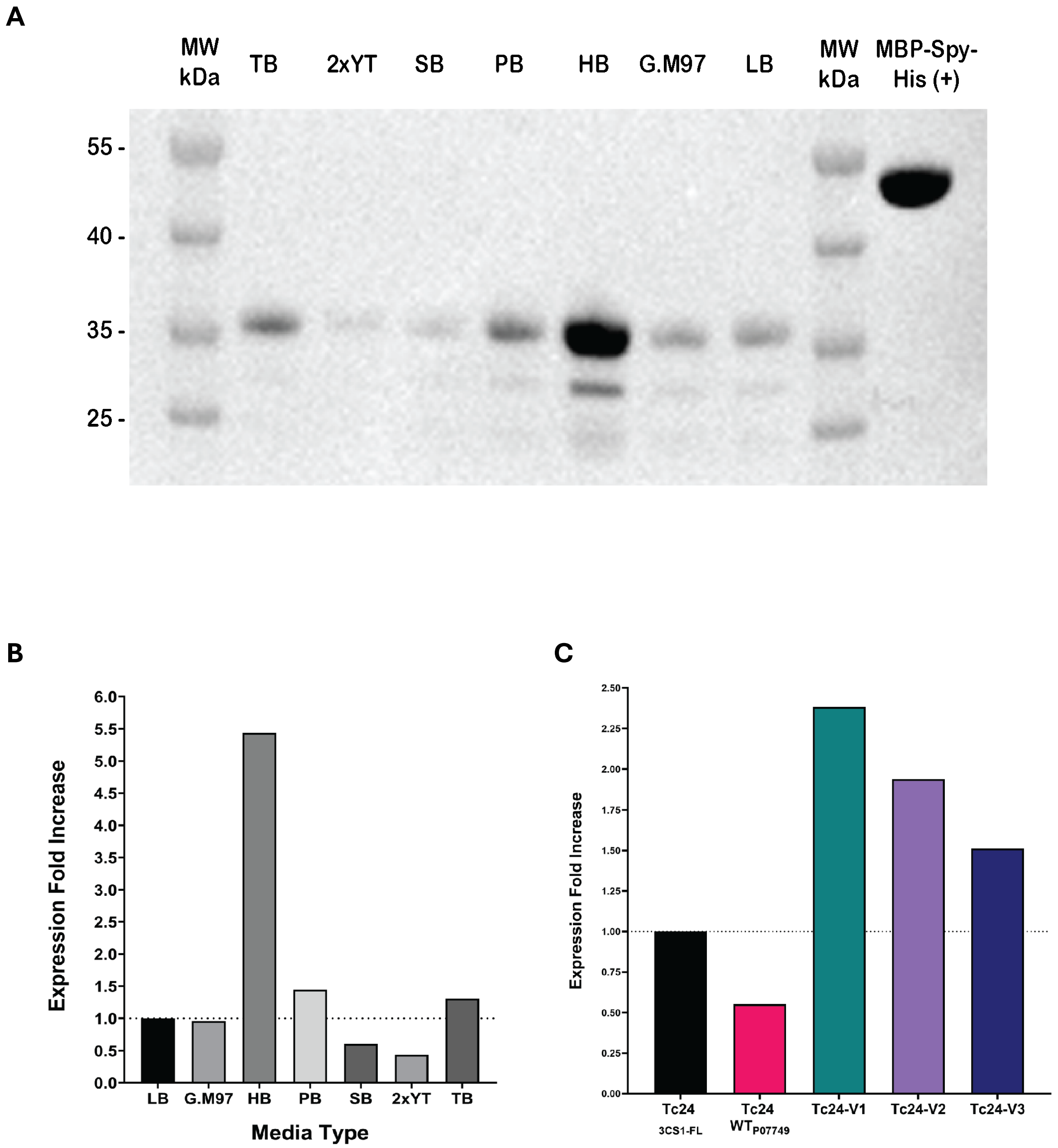


**Fig. S2** Media expression comparison of CL Brener Tc24. Soluble fraction Tc24 WT$_{Q4CTX0}$ Western Blot comparison from 100mL scale induction culture across Media Optimization Kit™ media types (**A**). Expression fold difference in densitometry analysis normalised against LB media, showing a 5.5 fold increase in Hyper Broth™ media (**B**). Quantified expression comparison between Tc24 WT constructs and Tc24variants (**C**).

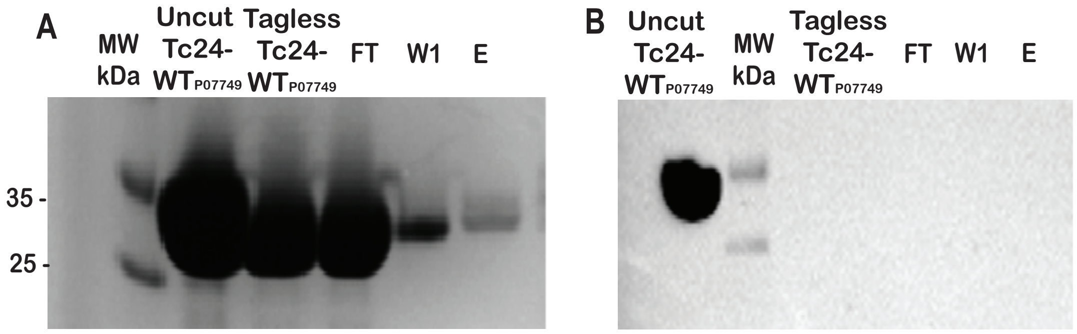


**Fig. S3** Tag cleavage from Tc24$_{3CS1\text{-}FL}$ (**A**) IMAC of Tagless Tc24$_{3CS1\text{-}FL}$ for uncut protein and tag removal post- overnight cleavage. (**B**) Western Blot for tag cleavage confirmation shown by presence and absence of signal in IMAC purification.

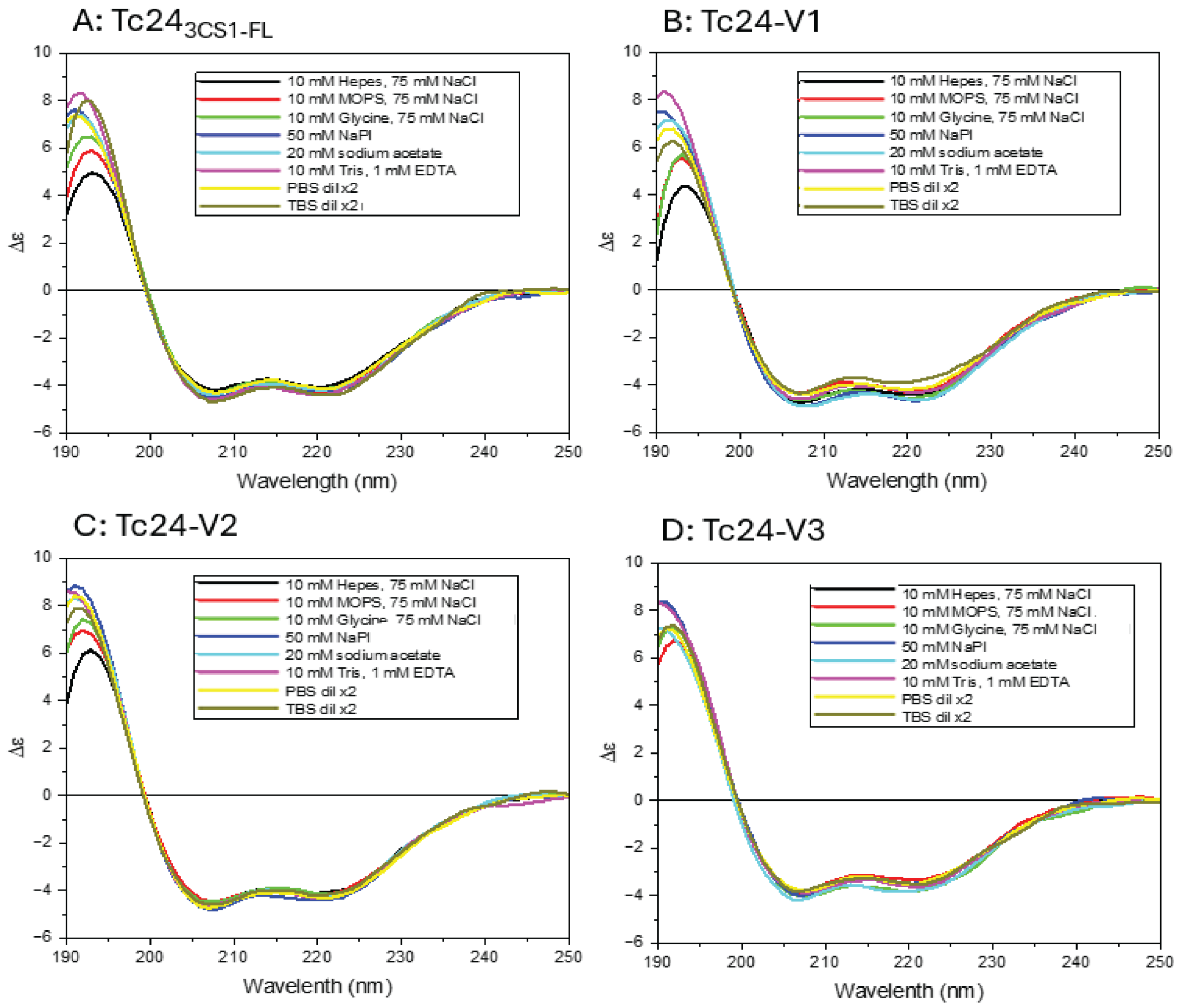


**Fig. S4** Far-UV (190-250nm) CD measurement of variants: (**A**) Tc24$_{3CS1-FL}$, (**B**) Tc24-V1, (**C**) Tc24-V2, and (**D**) Tc24-V3, using a nitrogen-flushed Module A end-station spectrophotometer with HTCD sample handling unit with 3 x12 quartz nominal 0.02cm pathlength string cells, 1s integration time, 300mA ring current, 22°C at B23 Synchrotron Radiation CD Beamline at the Diamond Light Source, Oxfordshire, UK. The initial observations show changes in secondary structure elements when exposed to various common formulation buffer components.

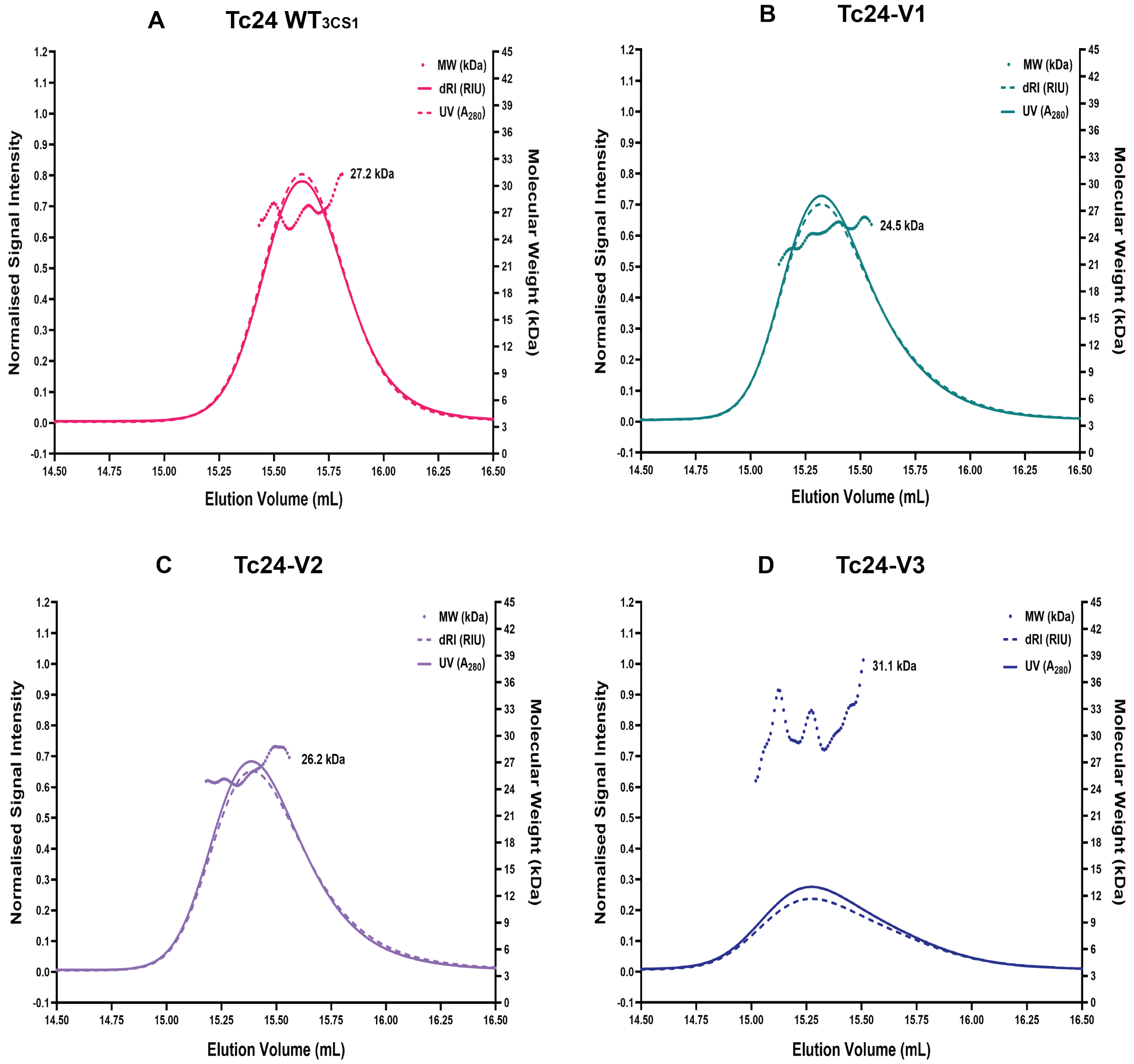


**Fig. S5 SEC-MALS analysis of lyophilised Tc24 Variants.** The UV $A_{280}$ (UV) data and differential refractive index (dRI) data recorded for each of the Tc24 constructs is shown. The molecular weight (MW, kDa) distribution across each peak is also indicated and, in all cases, this is consistent with each construct behaving as a monomer when resuspended in solution from lyophilised form. Tc24-$WT_{3CS1}$ (**A**) and variants Tc24-V1 (**B**, green), Tc24-V2 (**C**, violet), Tc24-V3 (D, blue).
